# Dityrosine photocrosslinking of native collagen bioinks for controlled shape-fidelity of bioprinted cardiac tissue constructs: probing the interplay between fibrillogenesis and covalent bond formation

**DOI:** 10.64898/2026.01.13.699021

**Authors:** Ana Nunes, Ankita Pramanick, Daniel Kelly, Rashmi Ramakrishnan, Vasileios Sergis, Vinh Khanh Doan, Hien Anh Tran, Khoon S Lim, Henrique Almeida, Andrew Daly

## Abstract

Collagen bioinks are widely used in biofabrication, but their relatively soft mechanical properties can lead to structural instabilities under cell-generated contraction forces. While synthetic functional groups can be conjugated for covalent crosslinking, these methods often disrupt natural protein fibrillogenesis, thereby compromising collagen fibre architecture. This work presents a strategy for the direct covalent stabilisation of native collagen bioinks with dityrosine bonds via visible-light photocrosslinking with ruthenium (Ru) and sodium persulfate (SPS), avoiding the need for polymer pre-functionalisation. Multimodal characterisation, including high-resolution microscopy, spectroscopy, mass spectrometry, and nanoindentation, identified photocrosslinking conditions that enhance collagen fibrillogenesis and reduce off-target polymer oxidation. Interestingly, the biofabrication process itself affected ultimate collagen fibre architecture, with shear-induced alignment during extrusion enhancing fibril proximity and self-assembly, overcoming inhibitory effects the crosslinkers had on fibrillogenesis via ionic and electrostatic interactions. Leveraging these insights, embedded bioprinting was used to fabricate cardiac constructs with high cell viability (>80%), where dityrosine crosslinking could be tuned to modulate geometric shape changes under cell-generated forces (1-15% shrinkage). Finally, the platform was used to bioprint anatomically accurate double-ventricle human heart models with robust shape fidelity. This research establishes a versatile photocrosslinking framework for bioprinting cardiac constructs with tunable shape stability using native collagen bioinks.

## 1. Introduction

Extrusion bioprinting enables the layer-by-layer assembly of 3D constructs that recapitulate the anatomical geometry and multicellularity of native tissues. In cardiac tissue bioprinting, collagen type I is commonly used for bioink synthesis due to its natural abundance in the myocardium ^1–3^. Physical crosslinking is the most widely used crosslinking strategy, where collagen fibrils self-assemble into fibrous networks following neutralisation at 37 °C ^4^. However, the chemical interactions formed during physical crosslinking are weak and reversible, leading to constructs with low mechanical strength that are prone to cell-mediated contraction and geometric instabilities ^1,5,6^. While we have shown that cell-driven shrinkage or shape-morphing, and the associated internal stresses, can promote the functional maturation of iPSC-cardiomyocytes ^6^, bioinks with tunable physical properties that can modulate the extent of structural remodelling will be required to ensure the production of cardiac tissues with predictable and consistent anatomical dimensions. Chemical crosslinking of collagen can be achieved using glutaraldehyde and carbodiimide crosslinkers ^7,8^ or by functionalising the polymer with methacrylate or norbornene side groups that support photocrosslinking reactions ^9,10^. While these approaches provide tunable control over mechanical properties, they still present various disadvantages regarding reaction complexity, bioproduct toxicity, and detrimental effects on cells ^11,12^. Furthermore, covalent crosslinking using synthetic functional groups can disrupt the protein’s ability to undergo natural fibrillogenesis, thereby compromising the resulting collagen fibre architecture ^13^.

Protein-based hydrogels containing tyrosine residues can also be photocrosslinked via a visible-light photoredox reaction involving ruthenium (Ru) - sodium persulfate (SPS), which generates covalent crosslinks between polymer chains through the formation of dityrosine bonds ^14^. This reaction enables photocrosslinking of collagen, gelatin, silk, and fibrin hydrogels in their native form ^15–17^, bypassing the need for pre-modification with functional groups such as methacrylates or norbornenes. Like other photocrosslinking reactions, a photocatalyst molecule is used, in this case, Ru, which harvests a photon and then transitions to an excited molecular form, Ru^2+*^. In this form, it acts as a reductant, donating an electron to the SPS molecules, inducing bond cleavage and generating sulphate anions and sulphate radicals (Figure 1a) ^18^. The resultant oxidative form of ruthenium (Ru^3+^) can oxidise tyrosine residues, generating tyrosyl groups that can react with nearby tyrosines to form dityrosine bonds (Figure 1a). The reaction is efficient and fast, and can support high cell viability due to the high absorbance of Ru in the visible light range, making it well-suited for bioprinting applications^15,19–21^. However, applying this system to collagen presents unique challenges due to its relatively low tyrosine content (approximately 0.5-1% ^22,23^) compared to more tyrosine-rich proteins like fibrin or silk. In one early application, FRESH bioprinting was used to create collagen type 1 conduits, which were then crosslinked in a bath of Ru/SPS ^24^. In another study, Ru/SPS was used to photocrosslink soft and stiff regions within collagen hydrogels to explore how stiffness gradients affect the contact guidance and durotaxis behaviour of human gingival fibroblasts ^25^. Despite the potential of Ru/SPS for photocrosslinking collagen bioinks in their native form, the fundamental mechanisms governing fibre network formation are poorly understood. Beyond the intended formation of dityrosine bonds, radicals generated during crosslinking, along with the ionic forms of the crosslinker molecules, will have multiple off-target molecular interactions that can alter the collagen monomeric structures and polymer network formation. For example, collagen fibrillogenesis is highly sensitive to the ionic environment and pH. Currently, few studies have examined how radical-based photocrosslinking methods, such as Ru/SPS, affect the composition of collagen or extracellular matrix (ECM)-based bioinks. In one example, using tyramine-modified hyaluronic acid and a riboflavin photoinitiator, it was demonstrated that reactive oxygen species (ROS) generated during photoinitiation can induce off-target polymer chain degradation ^26^.

**Figure 1.**
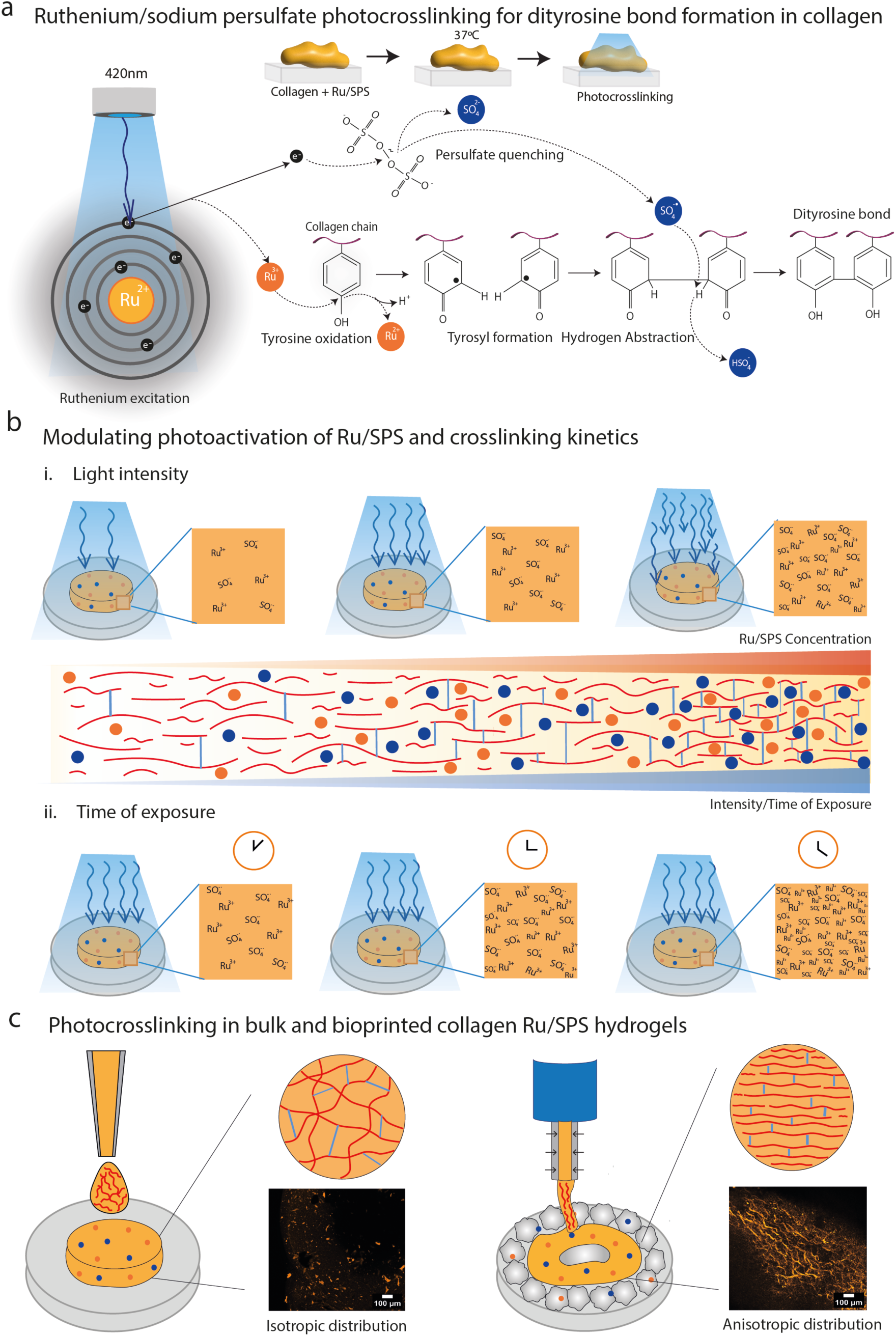
Ruthenium and Sodium Persulfate photocrosslinking system for dityrosine bond formation in collagen-hydrogels: a) Schematic representation of the Ru/SPS chemical reaction with tyrosine amino acids under blue light. b) Modulating Ru/SPS photoactivating by tuning variables like photocrosslinker concentration, light intensity (i) and time of exposure (ii), resulting in different dityrosine bond profiles. C) Demonstration of the collagen fibre distribution using conjugated collagen-FTIC. i) in-well bulk hydrogel and ii) printed hydrogel in a support bath.

In this study, we mechanistically investigated the Ru/SPS photocrosslinking system for native collagen bioinks for cardiac tissue bioprinting. Using a combination of spectroscopy, mass spectrometry, multiscale imaging, and nanoindentation analysis, we comprehensively characterised photoactivation kinetics and examined how off-target interactions between Ru/SPS molecules and collagen affect fibrillogenesis (Figure 1b). Building on these insights, we employed embedded bioprinting in granular supports to fabricate anatomically accurate human heart models using low-concentration collagen bioinks, demonstrating high cell viability with cardiac fibroblasts and induced pluripotent stem cell-derived cardiomyocytes (iPSC-CMs). Notably, our results revealed that the biofabrication process itself could dictate photocrosslinking outcomes, with shear-induced alignment during extrusion enhancing initial fibril proximity and fibrillogenesis, overcoming the effect of crosslinker ionic inhibition that reduced the impact of dityrosine bond formation in non-printed controls (Figure 1c). Using optimised dityrosine photocrosslinking conditions and low-concentration collagen bioinks, we then employed embedded bioprinting to fabricate cardiac tissue constructs with tunable mechanical properties capable of resisting contraction under cell-generated forces. These mechanistic findings establish a framework for leveraging Ru/SPS photochemistry to engineer mechanically tunable, structurally stable collagen bioinks in their native form without the need for chemical pre-modification.

## 2. Main

### 2.1 Dityrosine crosslinking of native collagen hydrogels using Ru/SPS photoinitiation

We began by exploring how Ru/SPS photoinitiation impacted dityrosine crosslinking in native collagen type 1 hydrogels (Figure 1a, b). A collagen concentration of 3mg/mL was selected to support cellular attachment and migration, considering that higher polymer concentrations typically result in denser networks that hinder cellular motility. This collagen solution was neutralised and mixed with different ratios of Ru/SPS (Figure 2a). After hydrogel solution preparation, the samples were physically crosslinked by incubation at 37°C for 30min, followed by photocrosslinking for 3mins using a 420nm light. Post-crosslinking, the samples had sufficient structural integrity to be easily scooped and transferred out from the plate wells (Figure 2a i). Lower SPS ratios (0.1 Ru / 0.25 SPS and 0.1 Ru / 0.50 SPS) showed better structural integrity than non-photocrosslinked controls, but still presented as soft gel-like structures (Figure 2a i). When the ratio of Ru/SPS was switched to 1:10 for the remaining formulations (0.1mM Ru / 1mM SPS, 0.3mM Ru / 3mM SPS and 0.5mM Ru / 5mM SPS), more stable hydrogels formed (Figure 2a i). These results support the importance of the sulphate radicals in the formation of dityrosine bonds that, in lower concentrations, are not sufficient to form a stable structure. Nanoindentation analysis revealed that photocrosslinking increased the hydrogels’ effective Young’s modulus from 200Pa (non-photocrosslinked) up to 500-800Pa, with significant increases observed for all Ru/SPS concentrations (Figure 2b iii). Interestingly, no significant differences were observed between 0.3mM Ru / 3mM SPS and 0.5mM Ru / 5mM SPS (Figure 2b iii). This data suggests that increasing the Ru/SPS concentration beyond 0.3mM/3mM will not increase the final stiffness of the hydrogel. This is a result of tyrosine saturation, also observed for other ECM-based hydrogels with Ru/SPS ^19,27^. It’s important to note that the tyrosine saturation concentration is dependent on polymer concentration, and higher Ru/SPS concentrations will likely affect stiffness when collagen concentration is increased. Interestingly, although lower SPS ratios (0.1 Ru / 0.25 SPS and 0.1 Ru / 0.50 SPS) appeared softer than other conditions (Figure 2a i), no significant differences in effective Young’s modulus were observed compared to the 0.1 Ru / 1 SPS condition (Figure 2b iii).

**Figure 2.**
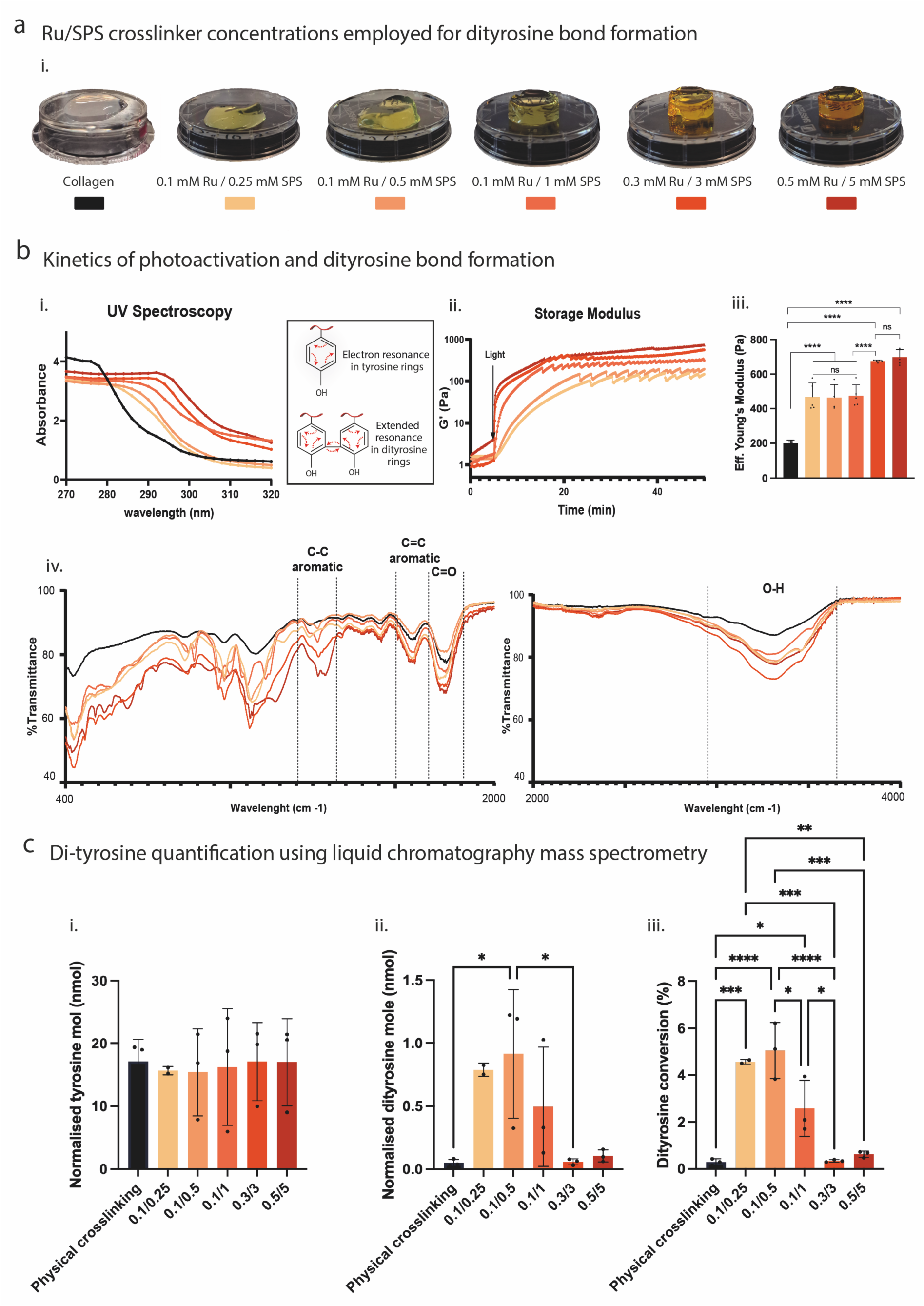
a) Collagen + Ru/SPS Hydrogels chemical and mechanical characterisation. i) Collagen + Ru/SPS hydrogels made with varied ratios of Ru/SPS concentration (0.1mM/0.25mM, 0.1mM/0.5mM, 0.1mM/1mM, 0.3mM/3mM and 0.5mM/5mM) compared to a collagen hydrogel. b) i) UV Spectroscopy of hydrogel samples for dityrosine bond formation qualitative analysis, demonstrating a spectrum shift as a result of resonance change between the aromatic rings of tyrosine after covalent bond formation. ii) Storage modulus analysis of the Ru/SPS hydrogels with varied Ru/SPS concentrations, demonstrating an increase in G’ as a result of photoactivation and crosslinking with light application after 5 minutes of the time-sweep test. iii) Resultant Effective Young’s modulus after photocrosslinking with one-way ANOVA with Tukey’s multiple comparison test (where **** denotes p < 0.0001 and ns denotes non-significant). iv) Fourier Transform Infrared Spectroscopy of the different Collagen + Ru/SPS bulk hydrogels after photocrosslinking and consequent freeze-drying, demonstrating the efficiency of the reaction by the C-C aromatic bonds and its off-target effects with the pronounced peak of C=O. C) i) Normalised total tyrosine content amongst varied photocrosslinker concentrations, showing no significant difference in content. ii) Normalised dityrosine content and iii) Degree of dityrosine conversion expressed as a percentage of dityrosine moles to total tyrosine moles were quantified by LC–MS. Data were normalised to the highest detected signal within each dataset, with statistical significance determined by one-way ANOVA with appropriate post-hoc multiple comparisons, where **** denotes p < 0.0001, *** denotes p < 0.001, ** denotes p < 0.01 and * denotes p < 0.05.

Complementing the nanoindentation analysis, we also performed rheological time-sweep analysis to assess how Ru/SPS concentration affected photocrosslinking kinetics (Figure 1b). These results demonstrated a correlation between the rate of polymerisation and Ru/SPS concentration (Figure 2b ii). Lower SPS concentrations (0.1 Ru / 0.25 SPS and 0.1 Ru / 0.50 SPS) demonstrated a slow increase of storage modulus, taking 18-20 mins to reach crosslinking saturation after light application. In contrast, formulations with 1:10 Ru/SPS showed a burst increase in the storage modulus, demonstrating the effectiveness of SPS molecules in rapidly generating dityrosine bonds. This indicates that even if higher Ru/SPS ratios (0.3/3 and 0.5/5) do not significantly change the final mechanical properties, the reaction proceeds faster because more Ru/SPS molecules are activated per unit time/area.

UV Spectroscopy measurements also demonstrated a correlation between Ru/SPS concentration and dityrosine formation, as evidenced by a continuous shift of the tyrosine peak at 275-280 nm to 290-300nm, with increased absorbance (Figure 2b i). This shift occurs due to electron delocalisation within the aromatic rings of tyrosine groups after covalent bond formation, altering their resonance profile. To complement this analysis, FTIR measurements were also performed to confirm the relationship between photocrosslinker concentration and covalent bond formation, as well as any possible side effects of the photocrosslinking reaction (Figure 2b iv). As confirmation of photocrosslinking efficiency and dityrosine bond formation, a band in 1200-1300cm^-1^ appears with decreased transmittance along with increased photocrosslinker concentrations. This band, not visible in collagen hydrogels, is associated with the C-C aromatic bonds, a dominant band that usually appears at 1400-1500 cm^-1^ ^29^ but can shift in the spectrum if changes in resonance along the aromatic rings occur. As mentioned before, the formation of covalent bonds between aromatic rings causes electron delocalisation, which alters the vibrational modes of CH bands, shifting the FTIR frequency band to 1200-1300 cm^-1^. As such, the FTIR demonstrated a direct correlation between dityrosine and photocrosslinker concentration.

A band at 3200-3400 cm^-1^ appeared with decreased transmittance for all collagen hydrogel compositions (Figure 2b iv). This is the result of physical crosslinking interactions, essentially hydrogen bridges involving hydroxyl groups. However, for photocrosslinked samples, the transmittance of this peak decreased (Figure 2b iv), suggesting that incorporating the Ru/SPS may inhibit the interactions required for fibril assembly and stabilisation into thicker fibres. Considering the ionic profile of the photocrosslinker molecules, we hypothesise that they might interact with the charged amino acids along the collagen chains. When incorporating these molecules before physical crosslinking, off-target interactions may occur, affecting the self-assembly of the fibres within the gel. Changes in fibril assembly can consequently impact the photocrosslinking system and covalent bond formation between small and weaker structures, which was further investigated in section 2.2. Additional side effects of the incorporation of Ru/SPS within the hydrogel solution are possibly demonstrated by the C=O band at 1600-1700 cm^-1^ (Figure 2b iv). Beyond electrostatic competition, these molecules can actively oxidise multiple amino acids with ketones and aldehydes, potentially increasing the formation of carbonyl groups. This oxidative damage can compromise the integrity of the collagen chain and the stability of the hydrogel network. These results highlight not only the efficiency of the photocrosslinking system in forming dityrosine bonds, already demonstrated for other natural polymers, but also demonstrate potential off-target effects.

To quantify tyrosine and dityrosine content in the fabricated collagen hydrogels, LC-MS measurement was performed. After normalisation with the highest detected signal, tyrosine content was comparable across all the formulations (both photocrosslinked and physically crosslinked control) with no significant difference observed (p>0.05). Meanwhile, the normalised dityrosine levels showed a strong dependence on Ru/SPS concentrations used to fabricate the hydrogels. In general, photocrosslinked hydrogels exhibited a clear increase in dityrosine content, showing significantly higher dityrosine formation at 0.1/0.5 mM Ru/SPS. However, further increasing the Ru/SPS concentration to 0.1/1 and 0.5/5 mM Ru/SPS resulted in a decline in dityrosine content, with no significant difference to the physically crosslinked control. Consistent with those trends, the degree of dityrosine conversion was the highest in the 0.1/0.5 mM Ru/SPS conditions and minimal in the physically crosslinked and high Ru/SPS levels (0.3/3 and 0.5/5). The results demonstrate that Ru/SPS initiated dityrosine formation peaks at intermediate crosslinking conditions, while excessive initiator concentrations impair dityrosine content and conversion rate.

Similar phenomena have been previously reported. The higher Ru/SPS concentrations (0.1/1 and 0.5/5 mM) were shown to significantly reduce the immobilisation of recombinantly expressed domain V of the human basement membrane proteoglycan perlecan (rDV) on silk scaffolds, as a result of diminished dityrosine coupling between tyrosine in rDV and silk ^30^. Another study, focused on photocrosslinked lipoaspirates with Ru/SPS, presented a similar trend to our study, in which an initial increase in dityrosine formation was observed at lower concentrations of Ru/SPS initiators (0.05/0.5 to 0.5/5 mM). The dityrosine content however decreased when higher levels of Ru/SPS (1/10 and 2/20 mM) were used ^31^. This reduction at high Ru/SPS concentrations is likely attributed to the high radical concentrations, which may promote radical quenching, chain termination, or over-oxidation, thereby limiting effective dityrosine coupling. Interestingly, we observed no significant increases in dityrosine content in the 0.3/3 and 0.5/5 mM conditions, even though these concentrations resulted in significant increases in mechanical properties (Figure 2b iii). This apparent discrepancy may be due to higher levels of Ru/SPS inducing alternative structural changes that contribute to hydrogel stiffening. For example, our FTIR analysis suggested off-target oxidative interactions, such as the formation of carbonyl groups, that could generate other non-specific crosslinking. The FTIR analysis also points to potential changes in collagen fibrillogenesis, which will affect the density and thickness of the collagen fibre network, thereby impacting construct mechanical properties independently of dityrosine bond formation.

Altogether, these results demonstrate that the Ru/SPS is an effective photoinitiation system for native collagen bioinks. Importantly, our results indicate that the crosslinker concentration needs to be optimised with respect to the total availability of tyrosine residues, and that increasing it beyond the saturation threshold can be detrimental, possibly interfering with fibrillogenesis. Having observed possible interference with fibril assembly in our spectroscopic analysis, the following section focused on imaging collagen fibres within photocrosslinked constructs and on investigating potential effects on long-term geometric stability.

### 2.2 Probing the impact of Ru/SPS photoinitiation on collagen fibre network structure

Building upon prior observations regarding the potential interference of Ru/SPS molecules with the collagen fibre assembly process, we next employed multiphoton and scanning electron microscopy (SEM) to evaluate the impact of photocrosslinking on collagen fibre structure. We also investigated whether the initial spatial distribution and positioning of collagen fibrils within a hydrogel volume could influence fibre assembly and the resultant network architecture under Ru/SPS-mediated photocrosslinking. Several variables can affect the final structure and architecture of the resultant collagen network. First, Ru and SPS are charged molecules that can electrostatically interact with other amino acids in the collagen chain. Such interactions may alter fibril assembly and network formation, which are primary determinants of hydrogel stability. Second, while the primary goal of the system is to stabilise the collagen chains via dityrosine bond formation, the sulphate radicals and photoactivated form of ruthenium have a high oxidative profile capable of altering amino acid chemistry and structure, as evidenced by our FTIR analysis. Furthermore, the relatively low tyrosine content in collagen (approximately 0.5-1% ^22,23^) and its accessibility within the protein’s quaternary structure may exacerbate non-specific Ru/SPS interactions.

Second harmonic generation (SHG) multiphoton microscopy demonstrated significantly enhanced collagen fibre formation in physically crosslinked collagen hydrogels (Figure 3a i). In contrast, the presence of 0.1mM Ru / 1mM SPS reduced fibre formation with minimal SHG signal observed, suggesting that electrostatic inhibition caused by the crosslinker molecules reduces fibrillogenesis into thicker collagen fibres (Figure 3a i). To further probe this outcome, we also conducted kinetic studies using UV Spectroscopy at different time points during physical crosslinking, analysing the collagen hydrogel’s turbidity over time, which can be used as a qualitative method to monitor crosslinking ^32^. We used four formulations, including unneutralised collagen, neutralised collagen, 0.1/1 Ru/SPS, and 0.5/5 Ru/SPS, which were kept at 37 °C for 40 minutes during analysis. As expected, the absorbance of the neutralised collagen solution increased over time due to fibrillogenesis and fibre assembly, stabilising after 30mins (Supplementary Figure 1a). Supporting the SHG analysis, constructs containing both Ru/SPS concentrations did not exhibit any spectral changes, maintaining continuous absorbance values (Supplementary Figure 1a). In a standard neutralised collagen solution, the protonation and deprotonation of amino acids occur, inducing chain electrostatic attraction, fibril proximity, and subsequent hydrogen bond formation. This results in the formation of a soft but compact network of thicker collagen fibres. When incorporating Ru/SPS during physical crosslinking, we hypothesise that electrostatic competition occurs, inhibiting natural attraction between collagen chains. The resultant photocrosslinked hydrogel will therefore be composed of smaller collagen fibrils that are linked with dityrosine bonds (Figure 3a ii). This inhibition of fibrillogenesis might also amplify polymer-solvent interactions, resulting in hydrogen bond formation between the polymer and water fraction of the hydrogel solution, generating a more soluble and unstable network structure (as previously observed by FTIR).

**Figure 3.**
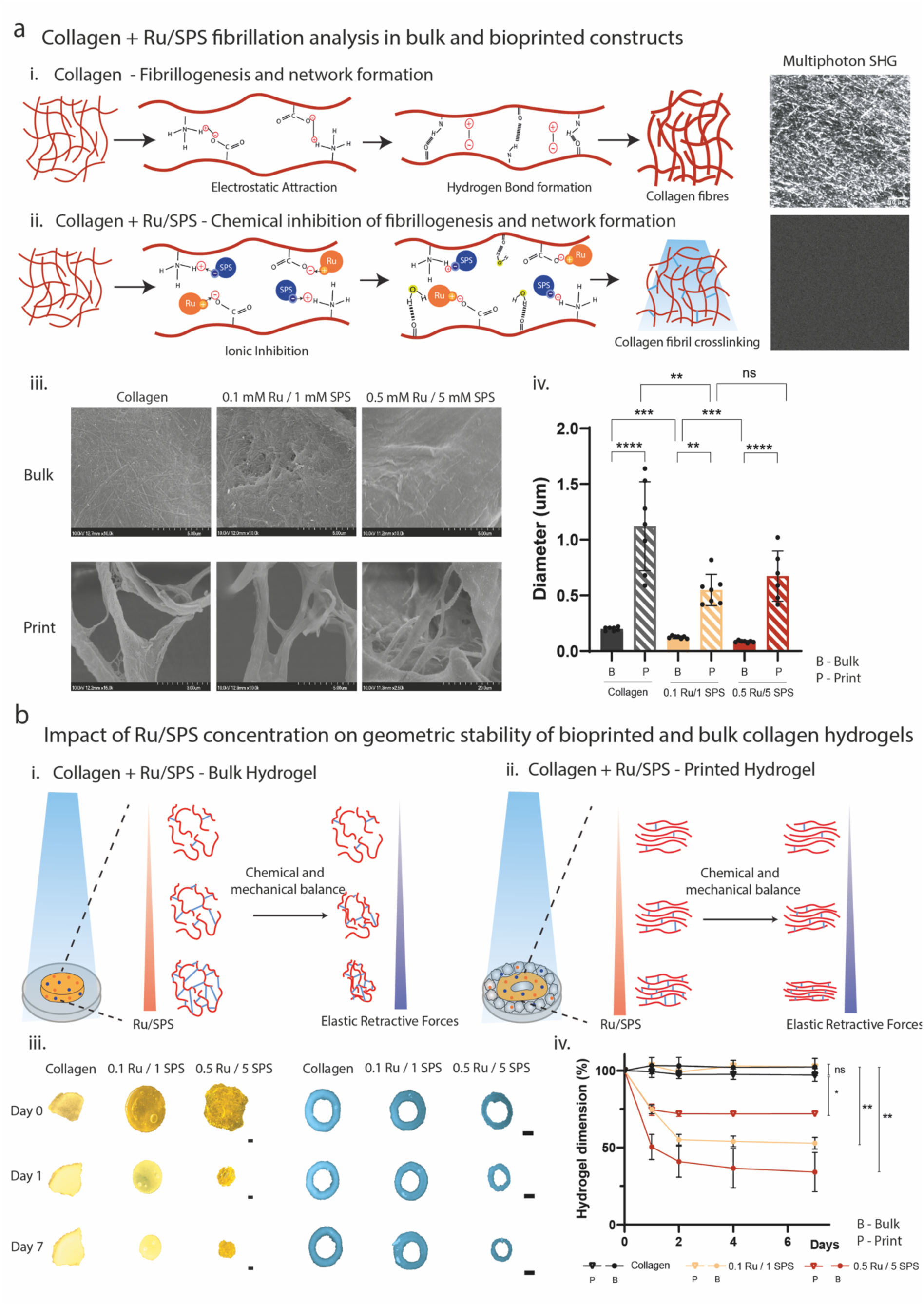
Collagen + Ru/SPS Hydrogels Stability studies. a) Schematic representation of the hypothesis behind Ru/SPS off-target interactions and self-assembly interference. i) Overview schematic of the chemical interactions (electrostatic interaction and hydrogen bonds) involved in collagen fibrillogenesis and respective Second Harmonic Generation (SHG) Microscopy image of the resultant fibre network distribution. ii) Hypothetic Ru/SPS off-target ionic interaction and decreased hydrogen bond formation, contributing to inefficient fibrillogenesis and network formation, incapable to be seen by SHG imaging. iii) Scanning Electron Microscopy of the 0.1mM/1mM and 0.5mm/5mM Ru/SPS bulk and printed gels compared to collagen hydrogels. iv) Resultant fibre diameter analysis demonstrating a significant decrease with the increase of photocrosslinker concentration, where **** denotes p < 0.0001, *** p < 0.001, ** p < 0.01. b) Schematic representation of the resultant hydrogel network stability after photocrosslinking Ru/SPS hydrogel samples. i) Overview representation of i) bulk Collagen + Ru/SPS hydrogel unstable network and resultant elastic retractive forces and ii) printed Col/Ru/SPS hydrogels where shear-forces might promote hydrogel network stability and reduced hydrogel contraction. iii and iv) Resultant hydrogel contraction profile (% of diameter change on day 0, 1, 2, 4 and 7 with n=3 replicates) of both bulk and printed Collagen + Ru/SPS hydrogels throughout 7 days (Scale bar 2mm), demonstrating significant contraction for bulk hydrogels, being significantly reduced in printed samples, where ** denotes p < 0.01 and * p<0.05.

Next, we assessed whether the biofabrication method itself could affect photocrosslinking outcomes. For example, shear forces during extrusion may enhance collagen chain nucleation and alignment. Under these shear-induced conditions, self-assembly and photocrosslinking will occur between more ordered and proximal fibrils. To do this, we used SEM to visualise collagen fibre thickness constructs prepared using embedded bioprinting of the collagen bioink into a granular support hydrogel (Figure 3a iii, iv). This was performed using both physical and photocrosslinking. As a control, we also analysed fibre thickness in constructs prepared using standard in-well crosslinking (termed bulk samples). Notably, collagen fibril/fibre thickness was significantly higher in bioprinted constructs for all crosslinking conditions, indicating that the shear forces during extrusion enhance the self-assembly process even with the Ru/SPS chemical environment (Figure 3a iii, iv). There was also a correlation between fibril/fibre thickness and photocrosslinker concentration for both bulk and bioprinted samples, with a decrease in thickness as Ru/SPS concentration increased (Figure 3a iii, iv). Spinning-disk confocal imaging of FITC-labelled collagen solutions also demonstrated that extrusion enhanced the visibility and thickness of collagen fibres compared with bulk crosslinking conditions (Figure 1c). These results further support the FTIR, UV spectroscopy, and SHG imaging results, indicating that the presence of Ru/SPS can impede early fibrillogenesis during physical crosslinking.

Interestingly, we also observed that the Ru/SPS concentration used for photocrosslinking strongly affected dimensional stability during long-term culture in PBS (Figure 3b). In bulk samples, 0.1/1 Ru/SPS resulted in a 50% dimensional (area) shrinkage over 7 days, with higher concentrations (0.5/5) generating a substantial 70% reduction in construct size (Figure 3b iv). These results suggest that Ru/SPS inhibition of physical crosslinking leads to deswelling of the constructs over time. To further probe these findings, we performed FTIR analysis on freeze-dried samples after seven days in PBS. We observed a pronounced band around 3200 cm^-1^, indicative of hydrogen-bond formation with water (Supplementary Figure 1b). In constructs photocrosslinked with 0.5/5 mM Ru/SPS, this band significantly decreased after 7 days, likely due to water expulsion and physical contraction of the hydrogel network (Supplementary Figure 1b). Similar trends were observed for the 0.1/1 mM Ru/SPS photocrosslinked constructs, although to a lesser extent. A pronounced band at 1650 cm^-1^ related to carbonyl groups was also identified, which we previously described as a potential oxidative effect of Ru/SPS entrapment in early stages. After 7 days, these bands also decreased, likely due to hydrogel contraction and reorganisation, promoting fibre interactions involving free OH groups. In contrast to their bulk form, constructs formed via embedded bioprinting in a granular support hydrogel were more geometrically stable over 7 days of culture (Figure 3b iii/iv). In particular, 0.1/1 mM Ru/SPS photocrosslinking conditions resulted in minimal shrinkage (Figure 3b iii/iv). The higher 0.5/5 mM Ru/SPS did result in a 25% shrinkage over 7 days, although this was still significantly less than in bulk controls (Figure 3b iii/iv). Similar to the FTIR analysis on bulk constructs, the peak at 3200 cm^-1^ decreased over 7 days, suggesting decreased interactions with water (Supplementary Figure 1b). Additionally, the carbonyl band presented a similar profile to the bulk hydrogels, suggesting that side-oxidation can also happen in bioprinted constructs, but due to network shrinkage, increases in collagen fibre proximity will promote interactions involving free OH groups.

The enhanced geometric stability in bioprinted constructs is likely due to the improved efficiency of the self-assembly process caused by shear forces during extrusion (Figure 1c), which will increase fibre thickness and proximity before dityrosine bond formation (Figure 3b ii). In contrast, bulk hydrogels prepared “in-well” exhibit lower self-assembly efficiency, resulting in networks of fine collagen fibrils that are crosslinked together by dityrosine bonds (Figure 3b i). The formation of covalent bonds between fibrils in these softer constructs may lead to network instability until chemical and mechanical balance is achieved (Figure 3b i).

Even if the photocrosslinking reaction occurs quickly after light activation, the polymer network formed must reach chemical, energetic, and mechanical equilibrium. The formation of covalent bonds between polymer chains increases polymer network density, generating elastic retractive forces that resist osmotic pressure driven by hydrophilic groups. As crosslinking density increases, the retractive forces increase, promoting hydrogel contraction and forcing a portion of the water out of the gel. The increased proximity of collagen fibres and the resultant dityrosine crosslinks formed in the bioprinted constructs are likely better able to resist shrinkage under these osmotic forces. Having explored how Ru/SPS concentration could affect the structural properties of bioprinted collagen hydrogels at multiple scales, we next examined how photocrosslinking conditions could affect the viability of encapsulated cells.

### 2.3 Evaluating the impact of Ru/SPS photocrosslinking on cell oxidative stress and survival

The previous results highlight how Ru/SPS photocrosslinking can impact the structural and mechanical properties of collagen hydrogels. Next, we evaluated how Ru/SPS reaction kinetics and the resulting hydrogel physical properties could influence cell survival and behaviour. We first investigated the toxicity profiles of the Ru and SPS crosslinkers themselves in physically crosslinked hydrogels (e.g., without light exposure), followed by evaluation after photocrosslinking. Simultaneously, we evaluated whether Ru/SPS-mediated stiffening could inhibit cell-mediated construct contraction.

Physically crosslinked control samples exhibited a concentration-dependent increase in toxicity as Ru/SPS concentration increased (Figure 4a ii and Supplementary Figure 2). Even in the absence of photoactivation, the crosslinker molecules can still interact with encapsulated cells, with lower cell viability observed in all conditions compared to Ru/SPS-free controls (Figure 4a ii). Additionally, partial exposure to ambient light may also trigger premature radical formation. These conditions induced intracellular oxidative stress, as evidenced by increasing DCFHDA fluorescence at higher Ru/SPS concentrations (Figure 4a i/iii). As expected, control non-photocrosslinked constructs underwent significant 50-60% contraction over 7 days of culture, with no significant differences between Ru/SPS concentrations (Figure 4a iv and Supplementary Figure 2). This phenomenon, which has been widely documented for soft, physically crosslinked collagen hydrogels ^5,6,33^, is driven by fibroblast-mediated contraction via actomyosin contractility.

**Figure 4.**
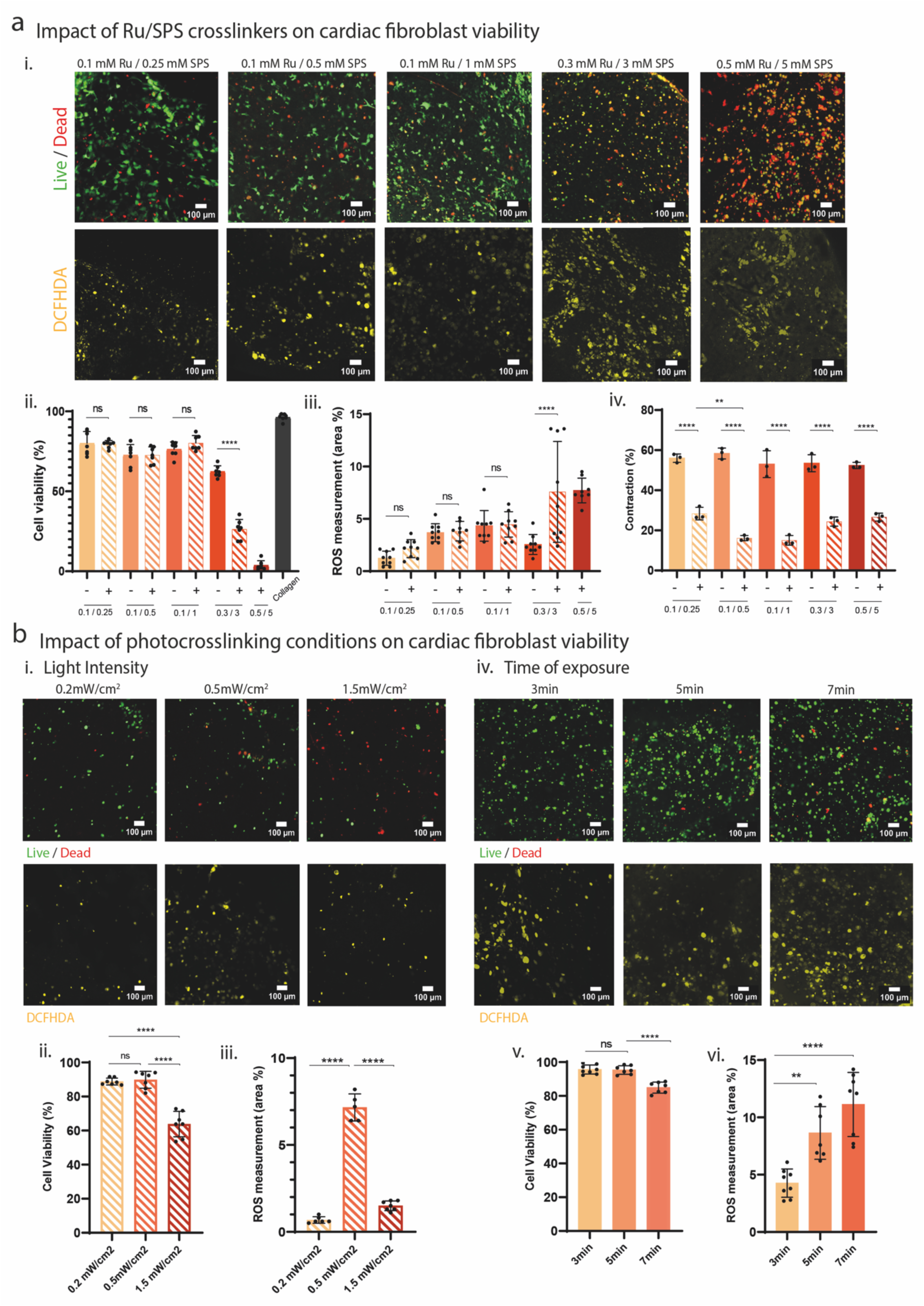
a) Influence of photocrosslinker concentration on Human Cardiac Fibroblasts and DCFDHA staining after 7 days of culture. i) Resultant cell viability quantification between non-photocrosslinked samples (Images on Supplementary Figure 2) and photocrosslinked samples (n=3 per formulation), demonstrating direct correlation between photocrosslinker concentration and cellular death rate using one-way ANOVA with Tukey’s multiple comparison test, where **** denotes p < 0.0001. ii) Reactive Oxygen Species staining by DCFHDA showing increased radical formation and cellular uptake with an increase of Ru/SPS concentration. iv) Quantification of hydrogel shrinkage (% of diameter change between day 0 and day 7 with n=3 replicates and one-way ANOVA with Tukey’s multiple comparison test, where **** denotes p < 0.0001 and ** p < 0.01) demonstrating reduced contraction in photocrosslinked samples compared to the control samples (Supplementary Figure 2). b) Impact of light variables in photocrosslinker activation and human cardiac fibroblast viability after 1 day of culture. i) Light intensity studies using 0.2, 0.5 and 1.5 mW/cm^2^ with resultant cell viability ii) and DCFHDA staining iii), demonstrating the detrimental effect of high intensity values. iv) Study of different times of exposure for photocrosslinking (3, 5 and 7 minutes) using 0.5 mW/cm^2^ with respective resultant cell viability v) and DCFHDA staining vi) suggesting that moderate light intensity with prolonged photocrosslinker activation can maintain cell viability.

Photocrosslinking significantly decreased contraction levels to 20-30% across all Ru/SPS conditions, demonstrating that dityrosine bond formation can resist cell-mediated shrinkage to generate geometrically stable constructs (Figure 4a iv and Supplementary Figure 2). For lower concentrations of Ru/SPS (0.1/0.25, 0.1/0.5, and 0.1/1), photocrosslinking resulted in cell viability levels of approximately 80% viability after 7 days, dropping to 20-30% when using higher concentrations (0.3/3, 0.5/5) (Figure 4a i, ii). DCFHDA staining demonstrated a gradual increase in intracellular reactive oxygen species with increasing photocrosslinker concentration (Figure 4a i, iii), demonstrating a relationship between Ru^3+^/SO^””^ generation and cellular toxicity. It should be noted that for all cell viability studies, 3mM of ascorbic acid was added to the media to balance the oxidative environment. As the Ru/SPS concentration of 0.1/1 mM was capable of resisting cell-mediated contraction while maintaining relatively high cell viability, it was selected for further photocrosslinking studies. Interestingly, we observed that cell encapsulation itself reduced construct shrinkage compared to cell-free controls in photocrosslinked constructs (i.e., shrinkage observed in Figure 3b). It is possible that cell interactions with the oxidative molecules reduced off-target effects. Additionally, the addition of ascorbic acid for the cell viability studies may have contributed to this effect.

To further optimise photocrosslinking conditions, we next assessed how light intensity and exposure time affected cell viability. For a fixed exposure time of 3mins, we observed high cell viabilities of 90% for light intensities of 0.2-0.5 mW/cm^2^, with viability dropping to 65% when the intensity was increased to 1.5 mW/cm^2^ (Figure 4b i and iv). Such results reflect the number of arriving photons per unit area, which can trigger rapid photoactivation and an acute oxidative environment within the hydrogel through excessive sulphate radical generation. Under these conditions, cells cannot respond quickly or activate endogenous antioxidant responses, triggering immediate cellular death pathways^34^. Cellular death can be triggered by different mechanisms of oxidative damage, such as membrane degradation and mitochondrial dysfunction, activating apoptotic and necrotic pathways. As a result of sulfate radical uptake by the cells, intracellular reactive oxygen species staining (DCFHDA) has also increased between 0.2 mW/cm^2^ and 0.5 mW/cm^2^, indicating increased radical generation with higher intensity. Interestingly, at the highest intensity of 1.5 mW/cm^2^, the DCFHDA fluorescence decreased significantly (Figure 4b i, iii). This is likely the result of more rapid cellular death and lower DCFHDA conversion within viable cells (Figure 4b i, iii) ^35^.

Next, we analysed how light exposure time impacted cell viability for the intermediate light intensity of 0.5 mW/cm^2^. In this case, prolonged exposure to moderate-intensity light will activate the photocrosslinkers at a controlled rate, gradually increasing the generation of Ru^3+^ / S*O*^-∂^_4_ (Figure 4b iv). Under these moderate conditions, the duration of exposure most likely affects cell viability by triggering cellular death pathways through the cumulative effect of Ru^3+^/ S*O*^-∂^_4_ and the gradual increase in oxidative stress. For exposure times of 3 and 5 mins, cell viability was high (>90%), with a significant decrease towards 80% when the exposure time was extended to 7 mins (Figure 4b iv, v). Although viability values remained unchanged between 3 and 5 mins of exposure, a significant increase in intracellular reactive oxygen species was observed between these conditions (Figure 4b iv, vi). This suggests that cells can sense and respond to moderate changes in the surrounding oxidative conditions (Figure 4b iv, v). Taken together, these results suggest that light intensity has a more pronounced effect on cell viability than exposure time, but also indicate that prolonged exposure times greater than 5 mins should be avoided. Based on our findings, photocrosslinking conditions with a light intensity of 0.5 mW/cm^2^ and an exposure time of 3 mins were employed for the bioprinting studies in the next section.

### 2.4 Embedded bioprinting of scaled-up, structurally stable heart constructs via dityrosine photocrosslinking of collagen bioinks

Next, we explored whether dityrosine photocrosslinking of collagen bioinks could be combined with embedded bioprinting methods to fabricate scaled-up, structurally stable heart constructs. By helping localise the bioink during extrusion and preventing subsequent structural collapse under gravitational forces, embedded bioprinting within a support hydrogel enables the fabrication of geometrically complex 3D constructs using low-viscosity collagen bioinks ^1,36,37^. Our previous research has demonstrated that embedded bioprinting of physically crosslinked collagen bioinks within granular supports can be used to create human heart models that undergo 4D shape-morphing via cell-generated forces ^6^. While internal stresses generated during 4D shape-morphing were found to enhance the maturation of iPSC-cardiomyocytes, these constructs were structurally unstable over long-term culture. With this in mind, we next explored whether Ru/SPS photochemistry could be used to induce dityrosine crosslinks to increase construct stiffness and modulate structural shape changes under cell-generated forces (Figure 5a). As per our previous research, we used an agarose-based granular support bath ^6^. While this material possesses suitable shear-thinning and self-healing properties for embedded bioprinting of collagen bioinks, it is also somewhat opaque, which, for photocrosslinking applications, can reduce the light penetration required for crosslinking (Figure 5a ii). We directly measured light penetration through increasing volumes of the agarose support, demonstrating attenuation of light intensity due to scattering of the light caused by the packed microgels (Figure 5a iii). This will decrease the final light intensity reaching the collagen bioink extruded into the support, reducing the number of photons that can excite ruthenium per unit area/time. For this reason, we adjusted the light-source distance from the sample to achieve higher intensity values close to 0.5 mW/cm^2^, which we had identified as optimal based on the results from section 2.3.

**Figure 5.**
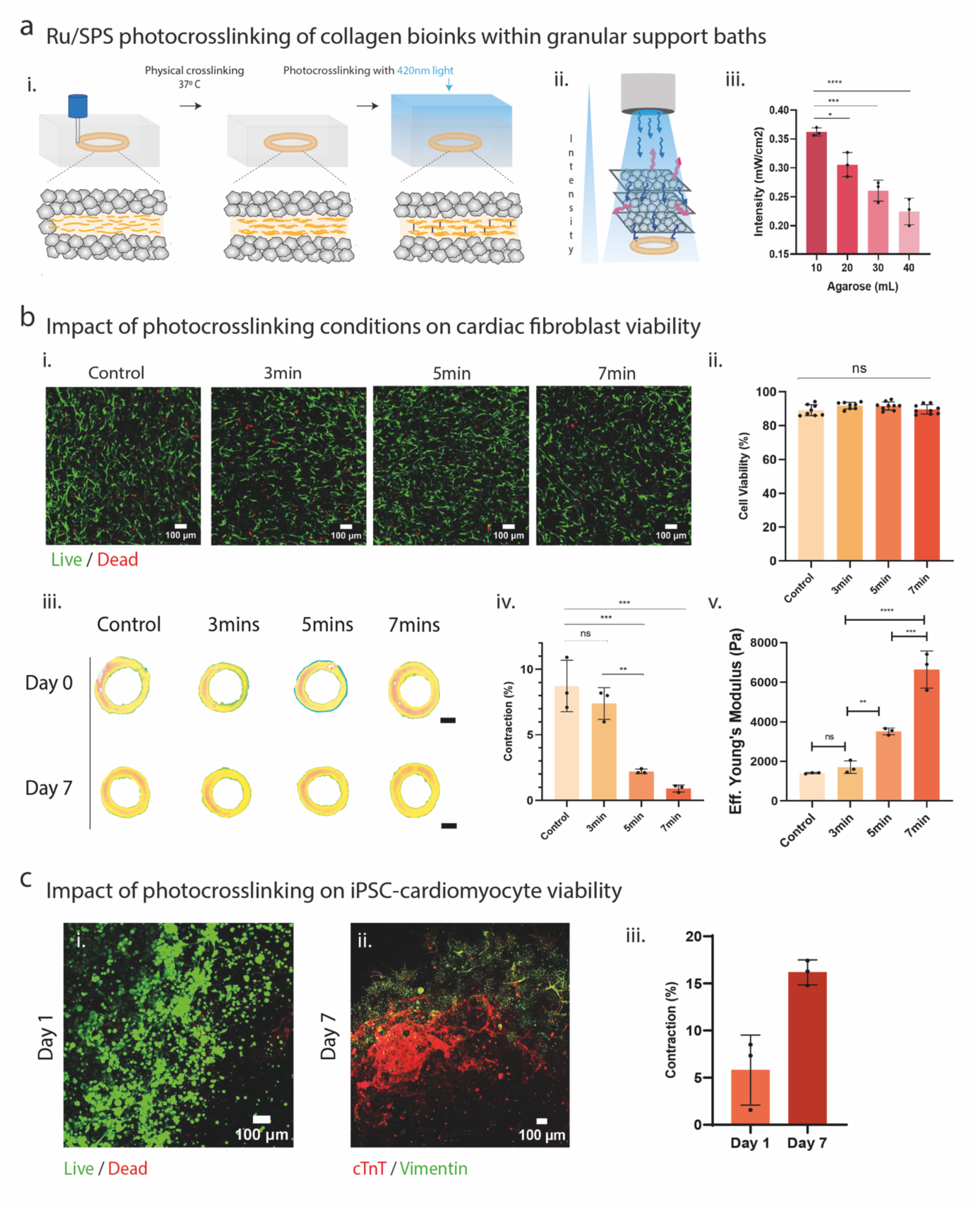
Collagen + Ru/SPS translation into an embedded bioprinting platform. a) i) Bioprinting Collagen + Ru/SPS printing in agarose support bath, followed by physical crosslinking and photocrosslinking using a 420nm blue light. ii) Light scattering events within the agarose support bath, resulting in reduced light intensity reaching the printed samples iii). b) Study the bioprinting of 0.1mM Ru / 1 mM SPS Collagen + Ru/SPS hydrogel with encapsulated cardiac fibroblast. i/ii) Quantification of cell viability of printed formulations with different times of exposure, with no significant difference between samples using a one-way ANOVA with Tukey’s multiple comparison test. iii/iv) Shrinkage analysis and quantification of the bioprinted samples (Scale bar 2mm), demonstrating direct correlation between time of exposure and hydrogel contraction. v) To support this shrinkage phenomenon, effective young’s modulus analysis was performed, demonstrating a significant increase with the time of photocrosslinking with blue light. c) Analysis of iPSC-cardiomyocyte printing viability with 0.1mM Ru / 1mM SPS Collagen + Ru/SPS hydrogel. i) Resultant cell viability after 1 day, demonstrating the biocompatibility of the system for iPSC-cardiomyocytes and ii) cardiac troponin (cTNT) and vimentin staining at day 7, demonstrating the cardiomyocyte phenotype retention. iii) Quantification of bioprinted samples shrinkage (% of diameter change between day 0 and day 7 with n=3 replicates), showing an increase in hydrogel contraction as a result of high cell density.

Considering this attenuation of photon arrival at the sample, we conducted viability and crosslinking analyses on collagen bioinks bioprinted within the support bath across increasing exposure times. Cardiac fibroblast cell viability remained high (85-90%) for all exposure times, with no significant differences compared to non-photocrosslinked controls (Figure 5b i, ii). Photocrosslinking also significantly reduced the structural shrinkage of the constructs relative to non-photocrosslinked controls, with lower shrinkage observed as light exposure time was extended (Figure 5b iii, iv). Nanoindentation analysis demonstrated that construct stiffness increased with longer photocrosslinking times (Figure 5b v), suggesting that the attenuation of shrinkage was likely due to increased collagen network stiffness, which could resist the effects of cell-generated contraction forces. These results demonstrate that Ru/SPS photochemistry can be used to modulate the mechanical properties of collagen bioinks bioprinted within support hydrogels. Notably, the mechanical properties of bioprinted constructs were stiffer compared to non-printed samples (Figure 2b iii). This may be due to the extrusion process, which enhances fibre proximity and alignment before photocrosslinking (Figure 1c, Figure 3a iii-iv).

The confocal images of cell viability also demonstrated that the cardiac fibroblasts displayed a spread morphology within the photocrosslinked constructs across all Ru/SPS concentrations (Figure 5b i). This native-like cardiac fibroblast morphology demonstrates that the introduction of dityrosine bonds can enhance construct stiffness and attenuate shrinkage without having detrimental effects on cellular behaviour. Building on this promising finding, we next encapsulated iPSC-cardiomyocytes within the bioink at a density of 20 million/ml in a co-culture with cardiac fibroblasts (7:3 ratio of iPSC-cardiomyocytes to fibroblasts). Following this, the embedded bioprinting platform was used to fabricate 5 mm diameter ring structures. Cell viability was high after 1 day of culture (Figure 5c i), and immunofluorescence staining for cardiac troponin T at day 7 demonstrated that the iPSC-cardiomyocyte population were able to retain their phenotype within the dityrosine crosslinked bioink (Figure 5c ii). The constructs were cultured for up to 14 days, and the emergence of spontaneous tissue contractions demonstrated that the encapsulated iPSC-cardiomyocytes were viable and functional within the dityrosine crosslinked collagen bioink (Supplementary video 1). It should be noted that the bioprinted cardiac tissue rings underwent significant shrinkage during culture, up to 15% by day 7 (Figure 5c iii). The increased shrinkage observed in these constructs is likely due to the higher cell densities employed (20 million/ml) compared to the fibroblast-only constructs (2 million/ml) (Figure 5b). For example, higher cell densities may alter crosslinking efficiency and will result in greater overall cell-generated contraction forces. It would be possible to modulate the extent of contraction for these higher-cell-density constructs by increasing bioink and crosslinker concentrations, as well as by altering photocrosslinking conditions.

Finally, we used embedded bioprinting and Ru/SPS photocrosslinking to fabricate scaled-up double ventricle heart models using collagen bioinks (Figure 6). To ensure the efficiency of the photocrosslinking reaction within a larger volume of support bath, the time and direction of light exposure were adjusted to maximise spatial efficiency, with the 420nm light source applied at multiple angles for 1-2 minutes, reaching a final exposure time of 8 minutes (Figure 6a). As per previous experiments, bioprinted constructs were physically crosslinked at 37°C for 30 mins before photocrosslinking. Physical crosslinking alone was insufficient to fabricate structurally stable constructs using the collagen bioink, with the constructs collapsing upon removal from the granular support bath (Figure 6b, - photocrosslinking). In contrast, the introduction of dityrosine bonds using photocrosslinking resulted in the formation of structurally stable constructs that exhibited robust shape retention (Figure 6b, + photocrosslinking), and cross-sectional imaging demonstrated that the constructs possessed open ventricular chambers (Figure 6c). These constructs were geometrically stable for at least 2 weeks in PBS and did not undergo degradation or shrinkage, highlighting the potential of the Ru/SPS photocrosslinking system for heart tissue bioprinting applications.

**Figure 6.**
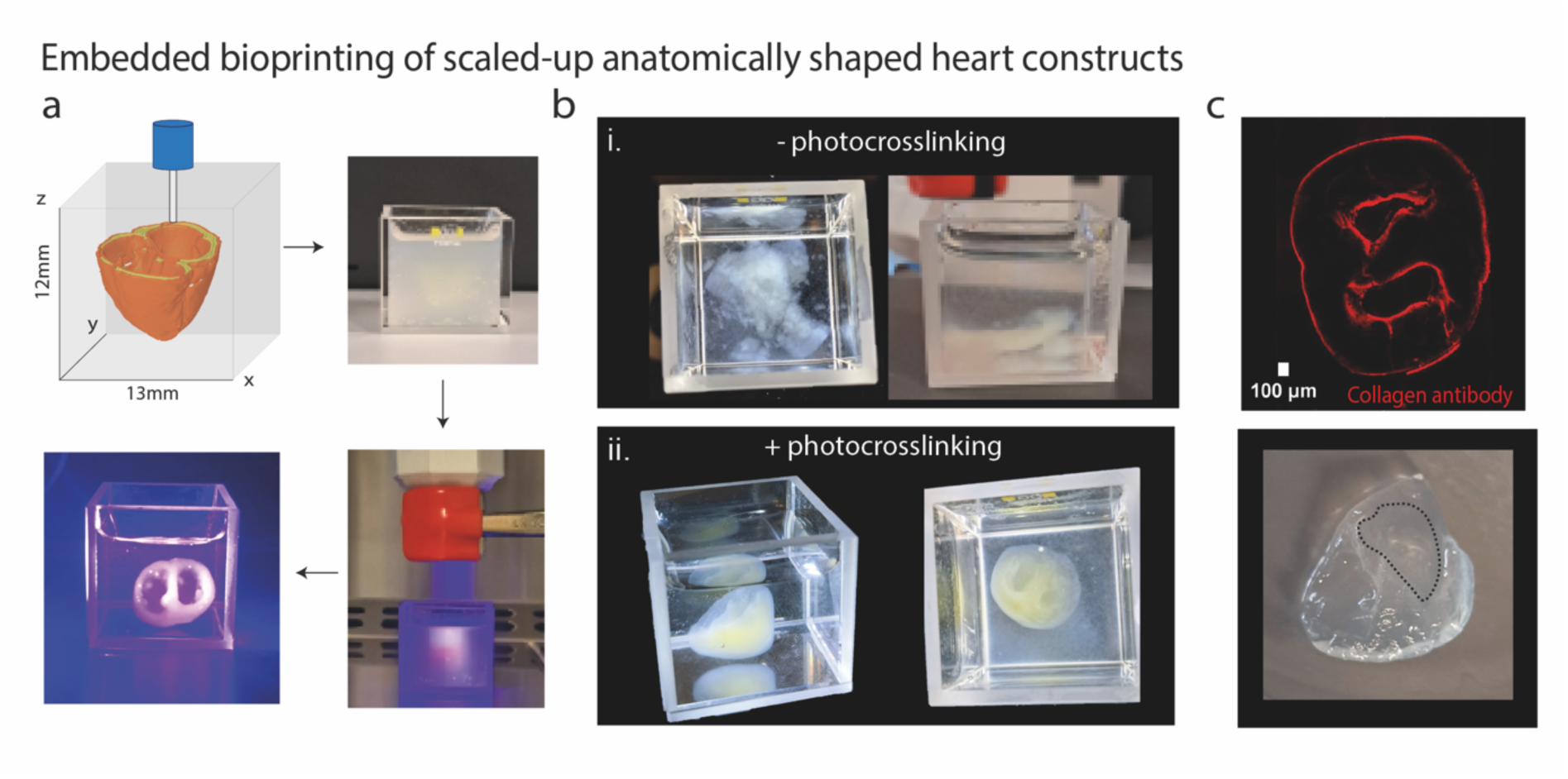
Embedded bioprinting of scaled-up structures using Collagen + Ru/SPS hydrogels. a) Representation of the heart cross-section g-code model, photocrosslinking procedure and support bath removal using 0.1mM Ru / 1mM SPS Collagen Bioink. b) i) Control printed sample (non-photocrosslinked) using Col_0.1mM Ru / 1mM SPS, demonstrating structural collapse after agarose support bath removal. ii) Photocrosslinked Col_0.1mM Ru / 1mM SPS samples showing structural resolution, printing fidelity and scaffold stability after support bath removal. c) Collagen type-I antibody staining of a slice of the printed cross-section structure, demonstrating cavity and shape retention.

Taken together, our results demonstrate that Ru/SPS photocrosslinking holds significant promise for heart tissue bioprinting applications. While collagen bioinks are widely employed for cardiac bioprinting due to their ability to support the long-term culture and maturation of encapsulated iPSC-cardiomyocytes ^2,6^, they remain susceptible to cell-mediated shrinkage and geometric instabilities. While internal stresses generated during shrinkage or shape-morphing can accelerate the maturation of iPSC-CMs ^6^, it is also important to develop tunable bioinks in which this shrinkage behaviour can be controlled to enable the bioprinting of cardiac tissues with predictable geometrical features. Cell-mediated shrinkage can be modulated by using higher-concentration physically crosslinked collagen bioinks ^1,6^, but physical crosslinking alone becomes insufficient to resist contraction, particularly when native-like cell densities and fibroblasts are employed. Recently, a shrinkage-resistant composite bioink was developed for cardiac applications using collagen and gallic acid-modified hyaluronic acid, in which the radical-generating capabilities of gallic acid, combined with riboflavin, were used to induce photocrosslinking ^5^. While this crosslinking mechanism reduced shrinkage and supported iPSC-cardiomyocyte maturation, larger ventricular constructs still underwent significant contraction, shrinking to 40% of their initial size after 12 days. In contrast, our dityrosine photocrosslinked collagen constructs exhibited enhanced geometric stability, with maximum shrinkage values restricted to 1-15% by day 7, depending on the degree of crosslinking and cell density. Future studies will explore the longer-term structural stability of constructs with native-like cellular densities (200-300 million/ml) to determine if dityrosine bonds can control anatomical fidelity under higher contractile loads.

### 2.5. Conclusion

In conclusion, this research establishes a robust photocrosslinking framework for bioprinting structurally stable, anatomically accurate cardiac tissues using native collagen bioinks and dityrosine bond formation. The primary advantage of the Ru/SPS system lies in its simplicity and the relative preservation of the native collagen protein structure compared to pre-conjugation approaches (e.g., collagen methacrylate). By leveraging native tyrosine residues inherent in the collagen network, this strategy eliminates the need for multi-component polymers, pre-functionalisation, or complex synthesis steps. Our results also provide unique mechanistic insights into how Ru/SPS molecules can initially impede collagen fibrillogenesis and electrostatic competition, and we also identified how shear-induced alignment during extrusion can attenuate this effect by enhancing initial fibril proximity. This synergy between the biofabrication method and subsequent dityrosine photocrosslinking reaction enabled the fabrication of structurally stable, anatomically accurate human heart models using extremely low collagen concentrations of 2.4 mg/ml. Furthermore, by optimising light intensity and exposure times, we were able to achieve high cell viability (>80%) for encapsulated cardiac fibroblasts and iPSC-cardiomyocytes in constructs in which mechanical stiffening could resist cell-mediated shrinkage to between 1-15%. Ultimately, this work offers a versatile and simple crosslinking strategy that expands the possibilities for engineering functional, scaled-up, and structurally stable cardiac tissues using native collagen bioinks.

## 3. Experimental Section

### Collagen Ru/SPS Hydrogel Synthesis

Ru and SPS stock solutions (Advanced Biomatrix #5248) of 50mM and 500mM were prepared in Phosphate Bovine Serum (PBS), stored at -20° C, respectively. Pure collagen bovine type I was purchased from Advanced Biomatrix (#5005, 3.1mg/mL) to synthesise the hydrogels by initially neutralising it using 10x PBS, 0.2M Sodium Hydroxide (NaOH) and deionised water, balancing its pH to 7.4, resulting in a final concentration of 2.4 mg/mL. To test the potential of the photoinitiator, different ratios of Ru/SPS were added to the collagen. After complete homogenization, the hydrogels were submitted to 30-40 minutes of physical crosslinking at 37°C and subsequently photocrosslinked using a 410-420nm blue light (TSLC N3535 1-chip UV LED, ILS). For the photocrosslinking step, different light intensities were tested by varying the distance between the light source and the sample, and different exposure times were also explored.

### UV-Spectroscopy

To monitor the formation of dityrosine, Collagen + Ru/SPS hydrogels were synthetized in 96 well plates (n=3 per formulation), incubated for physical crosslinking and submitted to 3 minutes of photocrosslinking. The UV-Vis spectra were recorded using a plate reader VARIOSKAN FLASH (Thermo Scientific) by running the samples from 200-800nm at room temperature and then analysed.

### Fourier Transform Infrared Spectroscopy

For chemical confirmation of dityrosine bond formation and to analyse other possible effects of the photo-reaction, the hydrogels were prepared as previously described and kept in -80°C overnight. After freezing, the hydrogels were lyophilised for 1 day and stored at -80°C for further analysis. Small pieces of the hydrogel were cut, analysed in ATR-FTIR Spectroscopy (SHIMADZU IRSpirit, A224157), and the data were exported for analysis.

### Rheological Analysis

To understand the efficiency and kinetics of the photocrosslinking process, time-sweep rheological tests were recorded using a Modular Compact Rheometer (MCR 302, Anton Paar) with a 25mm diameter probe with 1mm gap between sample at 1% constant strain.

### Nanoindentation Analysis

The effective Young’s modulus of the hydrogel formulations were obtained with an Optics11 Life Pavone Nanoindentor using a 29.5 μm tip radius and 4.18 N m^−1^ stiffness probe in air.

### Scanning Electron Microscopy (SEM)

To assess collagen fibre dimensions and resultant matrix of the different hydrogel formulations, the samples were synthesised, frozen and lyophilised for 1 day. After lyophilisation, samples were glued on double-sided carbon tape and sputter-coated with a 10 nm Au/Pd film. A Hitachi S-4700 EDX with an accelerating voltage of 10 kV was used to image the samples, and the images were analysed in Fiji software

### Multiphoton Microscopy

The collagen fibre structures were analysed using staining-free second harmonic generation (SHG) imaging using a multiphoton microscope (STELLARIS 8 DIVE). The excitation laser was tuned to 850-900nm to generate a SHG signal around 400-500nm. The same procedure was performed for a physically crosslinked collagen hydrogel and a photocrosslinked hydrogel (0.1 mM Ru / 1 mM SPS).

### Stability and Degradation studies

To investigate the stability of the photocrosslinked collagen hydrogels in both bulk (photocrosslinked in-well) and bioprinted forms as a function of Ru/SPS concentration, the following formulations were kept and analysed for 7 days: collagen hydrogel, Col_0.1mM Ru / 1mM SPS and Col_0.5mM Ru / 5mM SPS. To analyse the effect of the photocrosslinking system on hydrogel geometrical stability, multiple samples (n=3 per formulation) were freeze-dried and analysed under FTIR, on day 0 and day 7. In terms of hydrogel dimensional analysis, a camera system and software “Dinocapture 2.0” were used to record the size of each formulation (n=3) at different time points. The images were analysed on Fiji software, and the data were exported to GraphPad Prism 8.4.3.

### Primary Human Cardiac Fibroblast (HCF) Culture

Primary Human Cardiac Fibroblasts purchased from Promocell were cultured and expanded in Gibco MEM alpha-GlutaMAX medium (5% fetal bovine serum (FBS), 1% penicillin-streptomycin and 5 ng mL^−1^ Human Recombinant Fibroblast Growth factor-2 (FGF-2). For cell encapsulation in the hydrogels, cells were trypsinised with 0.5% trypsin-EDTA for 5 minutes and centrifuged at 300G for 5 minutes. For both cell viability trials and bioprinting experiments, the cells were used between passages 7 and 13.

### iPSC Differentiation to Cardiomyocyte and Encapsulation in Collagen + Ru/SPS Bioinks

Human Induced Pluripotent Stem cells (A18945, Fisher Scientific) were seeded on Corning Matrigel (354277, Fisher Scientific) coated six-well plates and expanded in Gibco Essential 8 medium (A2858501, Fisher Scientific) as previously described ^6^. For cardiomyocyte differentiation, iPSCs were cultured on Matrigel-coated 12-well plates, and differentiation was initiated when the cells reached 65-80% confluency using the GIWI protocol as previously described ^6^. For bioprinting experiments, the iPSC-cardiomyocytes were detached through incubation in Gibco™ TrypLE™ Select Enzyme 10X (A1217701, Fisher Scientific) for 20 mins, followed by centrifugation at 200G for 3mins. The iPSC-cardiomyocytes were then gently dissociated into single cells before being added to the collagen bioink. The cells were not purified but were simply trypsinised and mixed with cardiac fibroblasts (7:3 ratio of iPSC-cardiomyocytes to cardiac fibroblasts) at a final cell density of 20 million/ml. For the first 24h post-bioprinting, the ROCK inhibitor Y-27632 (72304, STEMCELL Technologies) (10µM) was added to the maintenance medium to promote cell survival. These bioinks were used to fabricate 5mm rings via embedded bioprinting in a granular agarose support hydrogel, and the constructs were then cultured within the support gel as previously described ^6^. The constructs were imaged using a Dinocapture camera to record their size and contractility.

### HCF Viability in Collagen + Ru/SPS Bioinks

After collagen neutralisation, the trypsinisation of confluent human cardiac fibroblast cultures was performed, resuspended in media (cell density 2 M cells/mL) and added to the neutralised collagen. Prior to a homogenization step, the photocrosslinker molecules were added, and 100μl of gel was placed in 96-well plates (n=3 per formulation). The gels were kept at 37°C for 30-40 minutes for physical crosslinking, shined with the 420nm blue light and detached from the walls of the wells with a needle. All of the Collagen + Ru/SPS hydrogels were kept in Gibco MEM alpha-GlutaMAX medium with 3mM of ascorbic acid. To analyse the impact of blue light intensity on cell viability, different intensities (0.2, 0.5, and 1.5 mW/cm^2^) and exposure times (3, 5, and 7 minutes) were tested.

### Live/Dead Staining

Live/dead staining was performed by treating samples with 2 μM Calcein-AM (Invitrogen, C3100MP) and 4 μM Ethidium homodimer-1 (Invitrogen E1169) in 1X PBS, followed by 40 mins incubation at 37°C. The samples were imaged using an Andor benchtop BC43 confocal microscope. To quantify cell viability, live and dead cells were counted (3 regions per sample) using Fiji software and the percentage viability was calculated using the formula below. This procedure was conducted for bulk hydrogel and subsequently for the 3D-bioprinted constructs.

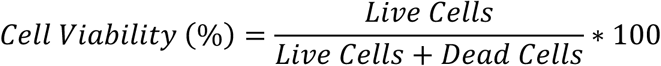

### Reactive Oxygen Species (ROS) – DCFHDA Staining

To analyse the intracellular reactive oxygen species, essentially radicals formed during the photocrosslinking, HCF-encapsulated hydrogels were stained with fluorescent redox probe (Diacetyldichlorofluorescein – DCFHDA) purchased from Abcam Cellular ROS kit (ab113851). The hydrogels were initially washed 2-3 times with 1X Buffer (10X Buffer diluted in ddH2O) and stained with a 20 μM DCFHDA solution (in 1X Buffer) for 45 minutes at 37°C in the dark. After incubation, the hydrogels were once again rinsed with 1X Buffer and analysed under a Confocal Microscope. The images were processed in Fiji Software by analysing the fluorescence per image obtained (n=3 and 3 regions per formulation).

### Agarose Support Bath Synthesis

To produce the support bath, agarose (Sigma-Aldrich) was used to prepare a 0.5 wt.% solution (in 1X PBS) and autoclaved at 121°C for approximately 1 hour. During cooling, the solution was stirred for 3 hours at 700 rpm. This procedure allows the generation of microgels via stir-generated mechanical forces as the agarose solution undergoes gelation upon gradual cooling to room temperature.

### Embedded Bioprinting for Collagen + Ru/SPS

High glass bottom μ-Dishes (35 mm, 81158, Ibidi) filled with 0.5 wt.% support bath was the bioprinted platform selected to pursue the different bioprinting analysis. Autodesk Inventor was used to create the 3D models for bioprinting (ring and heart cross-section), which were then converted to G-code by the Prusa Slicer software. The bioprinting experiments were performed using a custom in-house engineered Bioframe Bioprinter that employs a Puredyne extrusion printhead ^38,39^. The bioprinter was placed inside a biosafety hood to ensure sterile conditions. The collagen bioink temperature was maintained at 4°C during extrusion, controlled by the Puredyne cooling system around the print head. The bioinks were extruded into the agarose support bath using an embedded bioprinting method for all experiments. After bioprinting, all structures were physically crosslinked and consequently photocrosslinked. Finally, the bioprints were maintained at 37°C and 5% CO2 with regular media changes. To record and analyse morphological changes of the bioprinted structures, a camera system and software “Dinocapture 2.0” were used to record the scaffolds’ structure over time and analysed using Fiji software. The same conditions were used to bioprint the larger double-ventricle constructs, with larger glass dishes used to house the granular support hydrogel for embedded bioprinting.

### Immunostaining

For cardiac troponin T and vimentin staining of bioprinted cardiac tissues containing iPSC-cardiomyocytes and cardiac fibroblasts, constructs were fixed with 10% formalin overnight at 4°C, followed by Triton-X 100 (0.2% in PBS with 2%BSA) for 1h. After washing, the samples were blocked with 2% BSA in PBS for 2h. The primary bodies were used at a 1:200 dilution in blocking buffer, and staining was performed overnight. Following this, the samples were rinsed 3 times, and secondary antibodies were added (1:200 in blocking buffer) for 2h. For confocal imaging of the double-ventricle constructs, the samples were permeabilised and blocked, followed by overnight incubation with a primary collagen I Monoclonal antibody (Invitrogen, MA126771) to stain the collagen fibres. After washing with PBS, an anti-mouse Alexa 594 secondary antibody was used for confocal imaging.

### Dityrosine detection and quantification

A liquid chromatography tandem mass spectrometry (LC-MS) method was employed to detect both tyrosine and dityrosine content within the collagen hydrogels. Stable isotope-labelled internal standards (tyrosine (^13^C_6_) and dityrosine (^13^C_18_, ^15^N_2_) were used as reference to correct experimental variations in sample preparation and recovery, matrix effects, and instrument analysis, and to enable quantification of tyrosine and dityrosine contents.

- *Sample Preparation and Hydrolysis*: Collagen hydrogels were prepared as described in the above section prior to lyophilization. The lysophilized samples (∼2.5 mg) were hydrolyzed with added internal standards (1,000 nmol tyrosine (^13^C_6_) and 1,000 nmol dityrosine (^13^C_18_, ^15^N_2_)) in 6M hydrochloric acid: trifluoroacetic acid (2:1) containing 1% w/v phenol under a purged nitrogen condition at 150 °C for 1h. To remove the acid, samples were cleaned up using a solid phase extraction cartridge (Strata C18-E (55 µm, 70 Å)). Elutes were collected in 80% (v/v) methanol before concentrated via an Eppendorf centrifuge concentrator (Eppendorf, USA) and reconstituted in 0.1% (v/v) formic acid prior to analysis.
- *LC-MS/MS Method*: Standards and samples were analyzed using an online solid phase extraction method with a 5500 QTrap mass spectrometer coupled to a Shimadzu Nexera Prominence LC. Reconstituted samples were placed in the autosampler tray at 5 °C before being injected into a pre-equilibrated Imtake Intrada Amino Acid column (150×3.0 mm) from Shimadzu using 30% 100 mM ammonium formate (Solvent A) and 70% acetonitrile containing 0.1% formic acid (Solvent B). Isocratic elution was run for 2 min, followed by a linear gradient to 100% Solvent A over 7 min. The column was then held at 100% Solvent A for 2 min and subsequently re-equilibrated to the initial condition (30% Solvent A and 70% Solvent B) for 6 min. The column oven temperature was controlled at 40 °C, and the flow rate was maintained at 0.25 mL min^−1^ throughout the run. Samples were analysed and the areas of dityrosine and tyrosine peaks were calculated using Skyline. Tyrosine and dityrosine contents were quantified by comparing analyte peak areas to those of their corresponding isotopically labelled internal standards. Quantification was achieved by fragmenting the [M+H] + precursor ion and monitoring positive-ion mode that resulted from the loss of CH₂O₂ (Table S1, Supporting Information). All the samples were performed with three technical replicates (injections) and three biological replicates per group.

### Statistical Analysis

Experimental data were analysed and processed using Microsoft Excel, and GraphPad Prism 10 Software was used to prepare all graphs. All graphs are presented as the mean ± standard deviation, and sample numbers are provided in each graph and figure legend. Statistical analysis was performed using GraphPad Prism 10 software. For biological studies (i.e., with cells), the reported n values denote biologically independent samples. Unpaired t-test, two-way or one-way ANOVA tests were used depending on the number of independent variables within the experiment, and Tukey’s multiple comparison test was used to compare differences between means. P values are represented as follows: ns denotes not significant, * p <0.05, ** p < 0.01, *** p < 0.001, **** p < 0.0001.

## Acknowledgements

This publication has emanated from research conducted with the financial support of the European Research Council (Grant number 101077900), and the EU Commission Recovery and Resilience Facility under the Research Ireland Future Digital Challenge Grant Number 22/NCF/FD/10991G. The research was also financially supported by Galway University Foundation through the O’Dea PhD Scholarship to AN. This publication has also emanated from research supported in part by a grant from Research Ireland and is co-funded under the European Regional Development Fund under Grant number 13/RC/2073_P2. We would like to acknowledge the University of Galway Light Microscopy Imaging Core and Dr Karolina Salciute for their support with Multiphoton and SEM microscopy. The authors are grateful for the support and assistance of Olena Kudina for helping with the Nanoindentation experiments, and are thankful for the contributions of Dr Juan Gomez, Dr Amir Abdo and Rishi Suri to assisting with the chemistry experiments. Finally, we would like to thank Dr Tomas Calmeiro for helping with Scanning Electron Microscopy images during early studies of the project.

## Data availability statement

All data supporting the results of this study will be made available at the time of journal publication.

## Table of Contents Image and Text

A dityrosine photocrosslinking framework enables the fabrication of structurally stable, anatomically accurate cardiac tissues using native collagen bioinks, where shear-induced alignment during extrusion enhances early fibrillogenesis and fibre proximity for dityrosine bond formation. The platform supports high cell viability and resists cell-mediated shrinkage, providing a versatile and simple photocrosslinking strategy for bioprinting structurally stable cardiac tissues using native collagen bioinks.

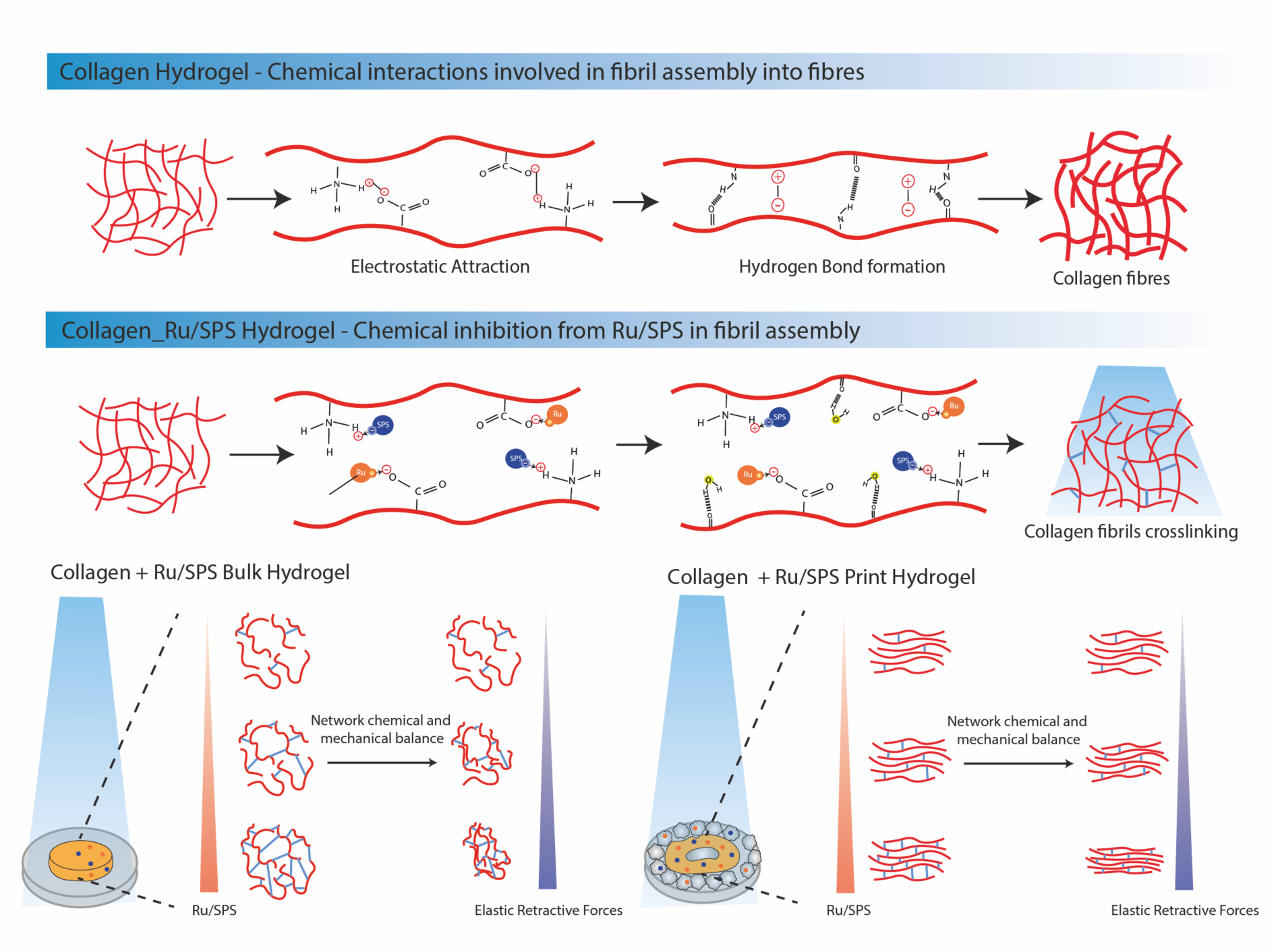

## Supporting Information S1

**Supplementary Figure 1:**
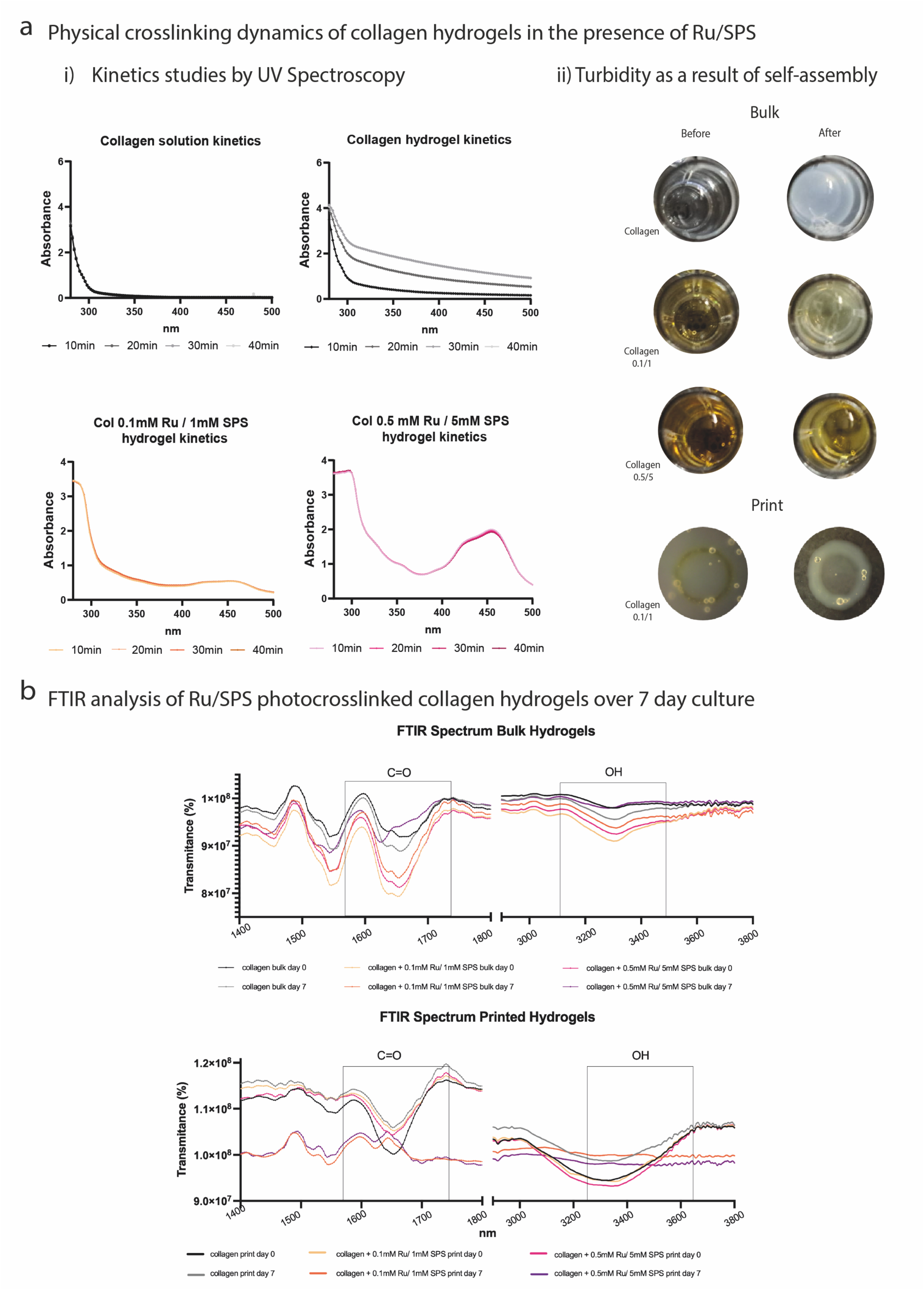
a) Studying the influence of Ru/SPS molecules in the fibrillogenesis of collagen by UV spectroscopy. I) Kinetic curve of collagen solution (pH=2.1-2.2), neutralised collagen (pH=7.2-7.4), Col_0.1mM Ru / 1mM SPS and Col_0.5mM Ru / 5mM SPS, demonstrating a gradual increase in absorbance of collagen hydrogel, as a result of self-assembly and decreased transmittance. ii) Illustration of the visual turbidity of the samples analysed, showing decreased turbidity with an increase in crosslinker concentration, matching the kinetic curves obtained. b) FTIR analysis of bulk and printed samples after synthesis and 7 days of culture in PBS, demonstrating reduction of the OH broad band around 3300cm^-1^, suggesting that hydrogel contraction over the days promotes post-interactions involving the OH groups.

**Supplementary Figure 2:**
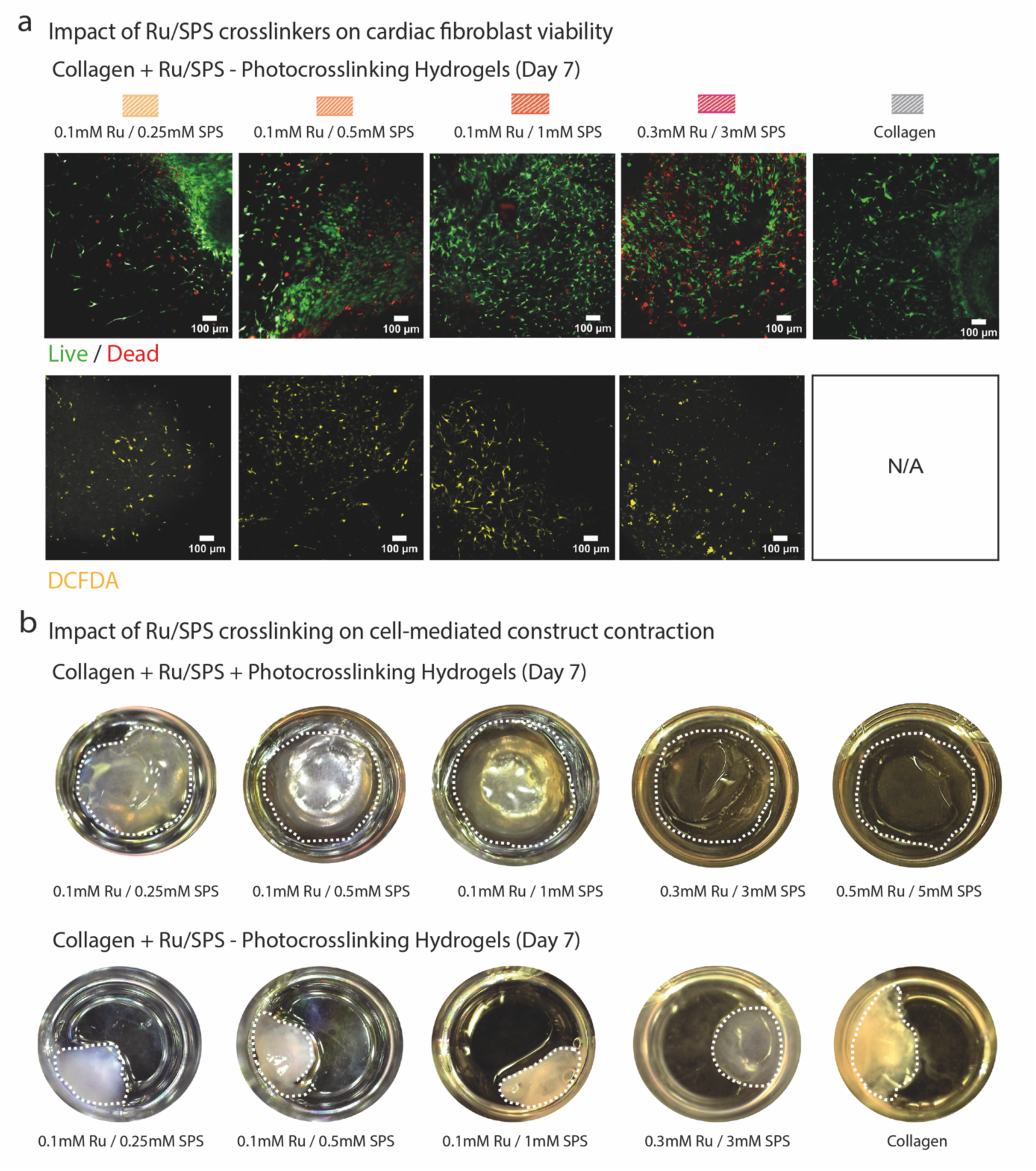
a) Influence of photocrosslinker concentration on control samples (non-photocrosslinked) with Human Cardiac Fibroblasts and DCFDHA staining after 7 days of culture, showing a direct correlation between photocrosslinker concentration and cellular death rate b) Hydrogel contraction of the varied Ru/SPS concentration (recorded with a Dinocapture system) of photocrosslinked (+ Photocrosslinking) and control samples (- Photocrosslinking) after 7 days in culture, demonstrating reduced contraction in photocrosslinked samples compared to the control samples.

**Table S1.** The mass-to-charge (m/z) ratio values of single-charged precursor and product ions, and the optimised parameters for quantification of each analyte in the LCMS measurement

| Analyte | Precursor (m/z) | Product (m/z) | DP | EP | CE | CXP |
| --- | --- | --- | --- | --- | --- | --- |
| Tyrosine (H<sup>+</sup>) | 182.02 | 136.09 | 45 | 10 | 19 | 12 |
| Tyrosine (<sup>13</sup>C<sub>6</sub>) (H<sup>+</sup>) | 188.06 | 142.12 | 35 | 10 | 18 | 7 |
| Dityrosine (H<sup>+</sup>) | 361.12 | 315.11 | 65 | 10 | 23 | 16 |
| Dityrosine (<sup>13</sup>C<sub>18</sub>, <sup>15</sup>N<sub>2</sub>) (H<sup>+</sup>) | 381.13 | 339.97 | 60 | 10 | 23 | 19 |

**DP:** Declustering Potential, **EP:** Entrance Potential, **CE:** Collision Energy, **CXP:** Cell Exit Potential

